# Nucleus accumbens neurons dynamically encode positive and aversive associative learning

**DOI:** 10.1101/2023.04.21.537763

**Authors:** Catarina Deseyve, Ana Verónica Domingues, Tawan T. A. Carvalho, Natacha Vieitas-Gaspar, Luísa Pinto, Nuno Sousa, Bárbara Coimbra, Ana João Rodrigues, Carina Soares-Cunha

**Affiliations:** Life and Health Sciences Research Institute (ICVS), School of Medicine, University of Minho, Braga, Portugal; ICVS/3B’s-PT Government Associate Laboratory, Braga/Guimaraes, Portugal; Clinical Academic Center-Braga (2CA)

**Author notes:** Both authors contributed equally (Catarina Deseyve, Ana Verónica Domingues). **Correspondence:** Ana João Rodrigues ICVS/School of Medicine University of Minho Campus de Gualtar 4710-057 Braga Portugal, **Email:**, &, Carina Soares-Cunha ICVS/School of Medicine University of Minho Campus de Gualtar 4710-057 Braga Portugal, **Email:**.

## Abstract

To survive, individuals must learn to associate cues in the environment with emotionally relevant outcomes. This association is partially mediated by the nucleus accumbens (NAc), a key brain region of the reward circuit that is mainly composed by GABAergic medium spiny neurons (MSNs), that express either dopamine receptor D1 or D2. Recent studies showed that both populations can drive reward and aversion, however, the activity of these neurons during appetitive and aversive Pavlovian conditioning remains to be determined.

Here, we investigated the relevance of D1- and D2-neurons in Pavlovian associations, by measuring calcium transients with fiber photometry during appetitive and aversive Pavlovian tasks. Sucrose was used as a positive unconditioned stimulus (US) and foot shock was used as a negative US. We show that during appetitive Pavlovian conditioning, D1- and D2-neurons exhibit a general decrease in activity in response to CS and to US across learning, with dynamic, and partially overlapping, activity responses to CS and US. During the aversive Pavlovian conditioning, D1- and D2-neurons showed an increase in the activity in response to the CS and to the US (shock).

Our data supports a synchronous role for D1- and D2-neurons in appetitive and aversion processing.

## Introduction

Daily, species learn and adjust their behavior according to their surroundings. Individuals attribute emotional value to otherwise neutral environmental cues if these are associated with a positive/rewarding or negative/aversive event (Cerri et al., 2014; Rescorla, 1994). These associations, in which a neutral stimulus becomes a conditioned stimulus (CS) once it is paired with an unconditioned stimulus (US) that is inherently rewarding (e.g. food, water or sexual stimuli) or aversive (e.g. shock) are called Pavlovian conditioning. The association transforms the neutral cue into a CS, that after learning has the ability to drive rewarding and aversive responses and contributes to learning (Day and Carelli, 2007; Pavlov, 2010).

Lesion and pharmacological evidence indicates that the nucleus accumbens (NAc) is critical for Pavlovian associative learning (Dalley et al., 2005; Di Ciano et al., 2001; Parkinson et al., 2002; Roitman et al., 2005; Setlow et al., 2003). In addition, seminal electrophysiological studies have shown that intraoral administration of sucrose leads to mainly inhibitory responses in NAc neurons, whereas administration the aversive compound quinine resulted in mainly excitatory responses (Roitman et al., 2005). Interestingly, as associations between CS and US occur, NAc neurons rapidly develop robust responses to the predicting cues (Roitman et al., 2005). Furthermore, it has been shown that NAc neurons increase activity during cue presentations but decrease during reward delivery (Ambroggi et al., 2011; Day et al., 2011; Gale et al., 2014). Similarly, in a Pavlovian conditioning approach task, the majority of recorded accumbal neurons exhibited a significant increase in firing rate during CS presentation, whereas the responses to reward were predominantly inhibitory (Day and Carelli, 2007; Wan and Peoples, 2006).

The NAc is mainly composed of GABAergic MSNs, which account for ∼95% of the accumbal neurons, being divided into those that express dopamine receptor D1 (D1-MSNs) or D2 (D2-MSNs) (Gerfen et al., 1990). D1-MSNs project directly to the ventral tegmental area (VTA), forming the direct pathway, while D2-MSNs, together with a sub-population of D1-MSNs, project to the ventral pallidum (VP), forming the indirect pathway (Kupchik et al., 2015; Soares-Cunha and Heinsbroek, 2023). Different studies have shown that both subpopulations are functionally divergent (Hikida et al., 2010; Kravitz et al., 2012; Lobo et al., 2010). However, more recent studies showed that optogenetic activation of either D1- or D2-neurons in the NAc supports self-stimulation (Cole et al., 2018), and can trigger positive and negative reinforcement (Namvar et al., 2019; Natsubori et al., 2017; Soares-Cunha et al., 2022, 2020, 2018, 2016).

Due to the similar properties between D1- and D2-MSNs in extracellular electrophysiological recordings, the genetic nature of neurons that respond to CS and US is not known. The development of genetically encoded calcium indicators opened new opportunities to identify these neurons. In the ventrolateral striatum, calcium elevations in D1- and D2-MSNs were observed at cue presentation in a motivation lever-pressing task (Natsubori et al., 2017). Plus, in a very recent study, putative NAc D1-MSNs presented multiphasic calcium events during CS presentation and reward delivery whereas D2-MSNs presented a monophasic event only after reward delivery (Skirzewski et al., 2022).

Although some studies point for a role of NAc neurons in cue-reward learning, how D1- and D2-MSNs encode positive or negative CS-US associations is far from being understood. Thus, to better understand the functional relevance of accumbal D1- or D2- neurons for the acquisition of CS-US Pavlovian learning, we used fiber photometry to register D1- and D2-neurons’ activity during positive and aversive Pavlovian conditioning in the same animals. We show that D1- and D2-neurons dynamically respond to both cue and reward in an appetitive Pavlovian task with sucrose as outcome. While D1- neurons show only an initial increase in activity in response to the cue, D2-neurons maintain an increased activity throughout learning. Interestingly, the activity of both neuronal populations decreases at reward exposure. In the aversive Pavlovian task, both neuronal populations increase activity in response to CS. D1-neurons present a more sustained increase in activity during foot shock in comparison to D2-neurons, that present sharper US response. These results show that both D1- and D2-neurons change activity during reward and aversion processing.

## Materials and Methods Animals

Male and female heterozygous D1-cre (line EY262, Gensat.org) and D2-cre (line ER44, Gensat.org) transgenic mouse lines (2-3 months of age) were used. All animals were maintained under standard laboratory conditions: an artificial 12h light/dark cycle with lights on from 8am to 8pm; with an ambient temperature of 21±1°C and a relative humidity of 50-60%. Mice were housed in type 2L home cages with a maximum of 6 mice per cage, with food (standard diet 4RF21, Mucedola, Italy) and water *ad libitum*, unless stated otherwise.

Behavioral experiments were performed during the light period of the light/dark cycle. All animals were kept divided according to gender and mouse strain from postnatal day 21. 5-10 minutes of handling was performed one week before behavioral experiments began, for 3 consecutive days, to avoid anxiety and stress responses. Animals were also habituated to all behavioral apparatuses for 3 consecutive days for 15 minutes before starting the behavioral tasks. Sample size used in behavioral tests was chosen according to previous studies; the investigator was not blind to the group allocation during behavioral performance.

All procedures involving mice were performed according to the guidelines for the welfare of laboratory mice as described in the European Union Directive 2010/63/EU. All protocols were approved by the Ethics Committee of the Life and Health Sciences Research Institute (ICVS) and by the national authority for animal experimentation, Direção-Geral de Alimentação e Veterinária (DGAV; approval reference #8332, dated of 2021-05-08). Health monitoring was carried out according to FELASA guidelines and all experimenters and animal facilities are accredited by DGAV.

### D1 and D2-cre line: mating and genotyping

The progeny produced by mating a D1-cre or D2-cre heterozygous transgenic male mouse with a wild-type C57/Bl6 female mouse was genotyped at weaning (postnatal day 21) by PCR.

DNA was isolated from tail biopsy using the Citogene DNA isolation kit (Citomed, Lisbon, Portugal). In a single PCR genotyping tube, the primers Drd1a F1 (5’-GCTATGGAGATGCTCCTGATGGAA-3’) and CreGS R1 (5’-CGGCAAACGGACAGAAGCATT-3’) were used to amplify the D1-cre transgene (340 bp), the primers Drd2 F1 (5’-GTGCGTCAGCATTTGGAGCA-3’) and CreGS R1 (5’-CGGCAAACGGACAGAAGCAT-3’) to amplify the D2-cre transgene (700 bp). An internal control gene (lipocalin 2, 500 bp) was used in the PCR (LCN_1 (5’-GTCCTTCTCACTTTGACAGAAGTCAGG-3’) and LCN_2 (5’-CACATCTCATGCTGCTCAGATAGCCAC-3’). Heterozygous mice were discriminated from the wild-type mice by the presence of two amplified DNA products corresponding to the transgene and the internal control gene. Gels were visualized with GEL DOC EZ imager (Bio-Rad, Hercules, CA, USA) and analyzed with the Image Lab 4.1 (Bio-Rad, Hercules, CA, USA).

### Surgery and optical fiber implantation

On the day of surgery, D1-cre and D2-cre mice (2-4 months old) were anaesthetized with sevoflurane (2-3% in oxygen), and their body temperature was maintained at 36-37°C using a closed-loop heating pad. An analgesic (buprenorphine) was administered before the beginning of the surgical procedure (0.05mgkg^-1^; Bupaq, RichterPharma, Austria). Animals were injected with a Cre-inducible AAV5-hSYN-FLEX-GCAMP6f-WPRE-SV40 (300nl, Addgene, Watertown, MA, USA) and AAV1-hSYN-FLEX-tdTomato (300nl (1:10 dilution), Addgene, Watertown, MA, USA) virus in the NAc (stereotaxic coordinates from bregma (Paxinos and Franklin, 2001): +1.3mm anteroposterior (AP), 0.9mm mediolateral (ML), and -4.5mm dorsoventral (DV)), using a Nanojet III injector (Drumond Scientific Company, Broomall, PA, USA), at a rate of 1nl/sec. After injection, the micropipette was left in place for 5min to allow viral diffusion.

After viral delivery, a fiber optic ferrule (400μm core, 0.50NA; Doric Lenses, Quebec, Canada) was implanted in the NAc using the injection coordinates (except of DV: -4.4) and was then secured to the skull with dental cement (C&B kit, Sun Medical, Shiga, Japan). At the end of the surgical procedure, mice were removed from the stereotaxic frame and postoperative care was carried out by administering analgesia (0.05mg kg^-1^ buprenorphine; Bupaq, RichterPharma, Austria) 6h post-procedure, as well as once every 24h during three successive days. A multivitamin supplement and saline were also administered post-procedure when necessary.

### Behavioral experiments

All animals performed the appetitive Pavlovian conditioning first and posteriorly the aversive Pavlovian conditioning.

#### Behavioral Apparatus

Behavioral sessions were performed in a custom-made operant chamber using pyControl software and hardware (17.8cm length x 19cm width x 23cm height) within a sound-attenuating box. In the appetitive Pavlovian conditioning, the chamber was composed by a central magazine, to provide access to 15μl of sucrose solution (20% wt/vol in water) delivered by a solenoid (for liquid dispenser), a cue-sound (70dB 5kHz), a house-light (100mA, 2.8W) installed on the top, and metallic floor. For the aversive Pavlovian conditioning, the chamber contained a house-light (100mA, 2.8W) installed on the top of the chamber, a cue-sound (80dB 2kHz) and a cue-light installed in one side wall and a grided floor with shocker. A computer was used to control the equipment and record the data, and a webcam (CMOS OV2710, ELP, Shenzhen, China) was used to acquire video.

#### Appetitive Pavlovian conditioning

(protocol adapted from (Patriarchi et al., 2018)). After 3 days of habituation to the behavioral box and the patch cable, mice (n_D1-cre_= 7, n_D2-cre_= 7) were exposed to 1-2 sessions of sucrose consumption, in which 15μl of a 20% sucrose solution were delivered every 30 seconds, for 30 minutes; this was done for animals to learn the position of the food port. After sucrose consumption session, mice started the appetitive Pavlovian conditioning in which a CS consisting of a 70dB 5kHz tone and a house-light (100mA, 2.8W) were turned on for 10 seconds; US of 15μl of 20% sucrose solution was made available at the 7^th^ second after CS onset. CS-US pairings were repeated 35 times per session, with a variable inter-trial interval (ITI) of 20-35 seconds (randomly assigned). Mice underwent a total of 12 sessions of appetitive Pavlovian conditioning. The behavior apparatus and the sucrose receptacle were disinfected with 10% ethanol between animals to remove any odor. For all sessions, nose poke data and photometry recordings were simultaneously obtained and synchronized through pyControl and pyPhotometry systems. To quantify CS-triggered behavior, number of nose pokes in the sucrose port were recorded during CS presentation; nose pokes were also registered during the ITI period.

#### Aversive Pavlovian conditioning

The same mice were familiarized to the behavior apparatus for 3 days for 10 minutes with the patch cable connected. Next, mice started a 3-day aversive Pavlovian conditioning protocol. All sessions started with 60 seconds of habituation period, with the house light on. The CS consisted of a 80dB, 2kHz tone plus a cue light. The US was a mild foot shock (0.5mA, over 1 second). During the first conditioning session, mice were exposed to 10 CS-US pairings. Each trial consisted of a random ITI (45-50 seconds) followed by a 10 second tone, which was immediately followed by electric foot shock delivered through the stainless-steel grid floor. On the second day (US omission), mice performed 30 trials in which 10 out of 30 tones (randomly assigned) were not followed by an electric foot shock. On the third day animals received a US extinction session, in which they received 30 trials with only CS exposure, but without US delivery (protocol adapted from (de Jong et al., 2019)). All sessions were recorded with webcams and photometry recordings were simultaneously obtained. To assess CS-triggered aversion, we measured freezing during the CS period from all sessions. The freezing response was defined as the time (seconds) that mice spent immobile (lack of any movement including sniffing) except respiration during the CS period and calculated as percentage of total cue time ((freezing time x 100)/cue duration).

### Fiber Photometry recordings

Recording of NAc D1- and D2-neuronal activity was performed during appetitive and aversive Pavlovian conditioning, using an open-source Python-based hardware and software for fiber photometry data acquisition - pyPhotometry (Akam and Walton, 2019). A fiber optic patch cable (1m long; 400μm core; 0.50NA; Doric Lenses, Quebec, Canada) was coupled to the implanted optic fiber with ceramic sleeve. The system used two LEDs (CLED_465; CLED_560; Doric Lenses, Quebec, Canada) via an LED driver circuit built into the system (with an adjustable 0-100mA output). Excitation light with two different wavelengths of 465nm for GCaMP6f (green) and 560nm for tdTomato (red), and an emission light were passed through a minicube (FMC5_E1(450–490)_F1(500–540) E2(550-580)_F2(600-680)_S; Doric Lenses, Quebec, Canada). GCaMP signal and a movement control signal from co-expression of tdTomato were recorded using the ‘2 color time-division’ acquisition setting. Light emitted by the indicators is transmitted back through the same optic fiber, splited from the illumination light and fluorescence from other indicators by optical filters and converted to electrical signals by two separate high sensitivity photoreceivers (Newport 2151 with lensed FC adapter; Doric Lenses, Quebec, Canada). Next, a Micropython Pyboard, with two analog and two digital inputs receive the fluorescence signals as analog voltage signals at a frequency of 130Hz (time division), which are then convert to digital signal via an RC lowpass filter. The digital inputs are read at the same sampling rate as the analog signals to enable *post hoc* synchronization of the photometry data with the behavioral hardware. The system uses one digital input to receive a TTL signal each time a behavioral event of interest (reward delivery or foot shock (or foot shock omission)) occurred.

### Fiber photometry data analysis

Initial analysis was performed in GuPPy, Guided Photometry Analysis in Python, a free and open-source package (Sherathiya et al., 2021). The first step was to remove artifacts. GuPPy uses a control channel (red) that includes smaller movement artifacts and photobleaching artifacts that are subtracted from the signal and used to normalize the data to determine changes from baseline fluorescence (ΔF/F). After establishing a control channel, a fitted control channel is used to calculate ΔF/F by subtracting the fitted control channel from the signal channel and dividing by the fitted control channel. After, a post-stimulus time histogram (PSTH) was calculated based on a defined window set around each event timestamp: for appetitive Pavlovian conditioning from -10 seconds to 20 seconds (CS onset at 0 seconds); for aversive Pavlovian conditioning from -10 seconds to 21 seconds (CS onset at 0 seconds).

After that, Python packages (Python version 3.9.12, Numpy version 1.21.5) were used to calculate the z-score from the data present on the PSTH, by subtracting the mean value of baseline activity (-10, -1) from all trials from each animal and dividing by the standard deviation.

The PSTH’s mean fluorescence activity (average of the z-score) and area under the curve (AUC; using Python package Scikit-learn version 1.0.2 (sklearn.metrics.auc)) in specific periods of time were then calculated. The intervals used for appetitive Pavlovian conditioning data were (time 0 seconds set at CS onset): pre-CS period (-2, 0); CS period: (0, 2); pre-US period (5, 7); US period (7, 9). For aversive Pavlovian conditioning data, the intervals were (time 0 seconds set at CS onset): pre-CS period (-2, 0); CS period: (0, 2); pre-US period (8, 10); US period (10, 12).

### Histology

#### Sacrifice and brain sectioning

At the end of all behavior procedures, mice were deeply anesthetized by a mixture of ketamine/medetomidine. Animals were then transcardially perfused with 0.9% saline, followed by 4% paraformaldehyde (PFA) solution. After, whole heads were immersed for 48h in 4% PFA so that the fiber track is clearly visible for histological analysis. Next, brains were extracted and then rinsed and stored in 30% sucrose at 4°C until sectioning. Sectioning was performed coronally, in 40μm slices, on a vibrating microtome (VT1000S, Leica, Germany) and slices were stored at 4°C on 12-well plates (or long-term storage in cryoprotectant solution at -20°C) until use. Slices from the area of interest (NAc) were selected using the Mouse Brain Atlas (Paxinos and Franklin, 2001).

#### Immunostaining (IHC)

To assess GCaMPf6 expression in D1-cre and D2-cre mice, brain slices containing NAc sections with fiber tracks were washed with phosphate buffered saline (PBS), and then permeabilizated with PBS-Triton 0.3% (PBS-T 0.3%). Blocking was performed for 1h using 5% Fetal Bovine Serum (FBS; Invitrogen, MA, USA) in PBS-T at RT. The primary antibody goat anti-GFP (1:750; ab6673, ABCAM, Cambridge, UK), was incubated overnight at 4°C with agitation, followed by PBS-T washes and subsequent incubation with the secondary fluorescent antibody Alexa Fluor® 488 donkey anti-goat (1:500; A11055, Invitrogen, Carlsbad, CA, USA). All antibodies were diluted in PBS-T with 2% FBS. Slices were washed with PBS-T, incubated with 4’,6-Diamidino-2-Phenylindole Dihydrochloride (DAPI, 1:1000; 62248, Thermo ScientificTM, Waltham, MA, USA) for nucleus staining, washed with PBS and mounted using Permafluor (Invitrogen, MA, USA). Slides were stored at 4°C and kept protected from light.

#### Image acquisition and analysis

Images from the NAc of D1-cre and D2-cre mice were collected in an inverted confocal microscope (Olympus FV3000, Tokyo, Japan). About 6-8 slices per animals were used for each analysis. Optic fiber placement was assessed to confirm if the activity detected was from the NAc region. For that, slices where the optic fiber was detected were classified according to the Mouse brain atlas (Paxinos and Franklin, 2001) to estimate the stereotaxic coordinates. Animals without viral expression or those with fiber placement outside the NAc were excluded from the analysis (2 D1-cre mice with fiber misplacement were excluded from the analyses).

### Statistical analysis

Statistical analysis was performed in GraphPad Prism 9.0 (GraphPad Software, Inc., La Jolla, CA, USA). Prior to any statistical comparison between groups, a Tukey test was performed to assess presence of outliers. Normality was assessed in all data analyzed by using the Kolmogorov-Smirnov (KS) test. Parametric tests were used whenever KS test > 0.05. If normality assumptions were not met, non-parametric analysis (Mann-Whitney or Wilcoxon test) was performed.

For behavior, two-way analysis of variance (ANOVA) for repeated measures was used to assess learning in D1- and D2-cre mice (factors used: nose pokes during CS *versus* nose pokes during ITI, across days of training), and to analyze percentage (%) of freezing throughout the trials on day 1. Bonferroni’s *post hoc* multiple comparison test was used for group differences determination. Additionally, a linear regression was also performed to assess the correlation between number of nose pokes during CS and days of training. Statistical analysis between two time-points was made using two-tailed paired Student’s t-test, to compare percentage of freezing in the first and last trials on day 1 and to compare percentage of freezing in the CS-US trials and CS-only trials on day 2. One-way ANOVA for repeated measures was performed to compare percentage of freezing in the first 5, intermediate 5 and last 5 trials of day 3 of the aversive Pavlovian conditioning.

Regarding fiber photometry data analysis, to determine whether activity of the PSTH (z-score) significantly (p<0.05) increased or decreased by a stimulus during the appetitive Pavlovian conditioning, an ANOVA was applied to every 2 seconds spanning from 2 seconds before CS to 6 seconds after CS, or from 2 seconds before US to 10 seconds after US. To determine whether activity of the PSTH significantly increased or decreased by a stimulus during the aversive Pavlovian conditioning an ANOVA was applied to every 2 seconds spanning from 2 seconds before CS to 9 seconds after the trial. Bonferroni’s *post hoc* multiple comparison was used for time-point differences determination. For simplification purposes, only significant differences detected between the 2 second period before and the 2 second period after event occurrence are represented in the graphs (Figures 2, 4). To assess differences in mean fluorescence from day 1, day 6 and day 12 of appetitive Pavlovian conditioning a One-way ANOVA was done; Bonferroni’s *post hoc* multiple comparison was used for group differences. To assess differences between AUC of D1- versus D2-neurons, a two-tailed Student’s t-test was done.

Data are presented as mean ± standard error of mean (SEM) or as min to max. Individual points are shown in the graphs. Statistical significance was accepted for p ≤ 0.05. All significant statistical comparisons are presented in the results section. All statistical analyses were run in GraphPad Prism 9.0 (GraphPad Software, Inc., La Jolla, CA, USA).

## Results

### Mice acquire proper responses to appetitive Pavlovian conditioning

To examine real-time activity of NAc D1- and D2-neurons *in vivo* during associative learning tasks, we injected D1-cre and D2-cre mice (n=7 per mouse line) in the NAc core with an AAV expressing a cre-dependent calcium indicator GCaMP6f, in combination with an AAV carrying a vector with tdTomato that was used for motion correction; calcium transients were recorded through an optic fiber (Figure 1A) in freely behaving mice. Viral expression (Figure 1B) and optic fiber placement (Supplementary Figure 1A-F) was confirmed for all animals.

**Figure 1.**
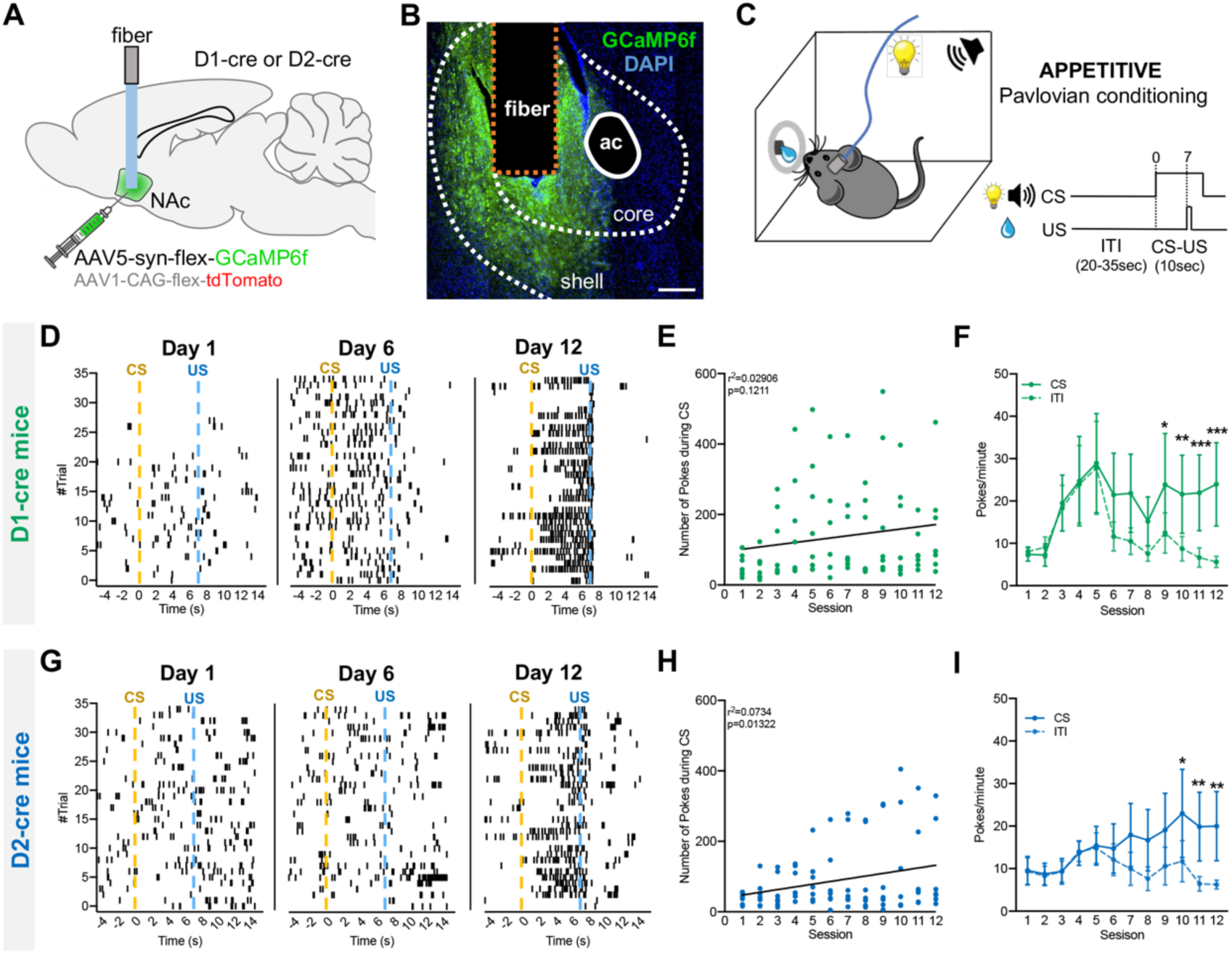
Acquisition of appetitive Pavlovian conditioning by D1- and D2-cre mice during fiber photometry recordings of NAc neurons. **A** Schematics showing the strategy for fiber photometry recording of GCaMP6f in NAc D1- or D2-neurons. **B** Expression of GCaMP6f in NAc and fiber tip location; scale bar: 250 μm. **C** Appetitive Pavlovian conditioning involved learning to associate a 5-kHz tone and house light (conditioned stimuli - CS) with 20% sucrose reward (unconditioned stimulus - US). Representative poke raster of a **D** D1-cre and a **G** D2-cre mouse, depicting individual pokes that were detected during day 1, day 6 and day 12 of behavioral learning. **E** Correlation between number of pokes and Pavlovian session in D1-cre mice, and in **H** D2-cre mice. **F** Number of pokes/minute during CS and ITI in D1-cre animals, and in **I** D2-cre mice. Data are means ± SEM. * p≤0.05, ** p ≤ 0.01, ***p ≤ 0.001.

Four to six weeks after surgery, mice performed the appetitive Pavlovian conditioning. After habituation to the chambers and to sucrose solution, mice were trained in the appetitive Pavlovian conditioning for 12 consecutive days. Each session involves 35 pairings of a CS (tone and house light) with the delivery of 15μl of a 20% sucrose solution as US (Figure 1C).

Initially, both D1- and D2-cre mice did not show robust anticipatory pokes in response to the cue, but, after multiple sessions of Pavlovian conditioning animals display consistent anticipatory poking, indicating that the cue-reward association had been learned (example of D1-cre mouse in Figure 1D; D2-cre mouse in Figure 1G). An increase in the number of pokes in the food receptacle during the CS period over days of training was observed in D1-cre, although not statistically due to high variability (r^2^=0.029, S=128.5, *p*=0.1211, Figure 1E); and D2-cre mice (r^2^=0.073, S=94.27, *p*=0.0131, Figure 1H). In addition, a significant higher number of pokes per minute were detected during the CS period in comparison with ITI (D1-cre: cue *vs* ITI, F_11,6_=3.0, *p*=0.0002, Figure 1F; D2-cre: cue *vs* ITI, F_11,6_=4.0, *p*=0.0046, Figure 1I).

### Temporal evolution of NAc D1- and D2-neurons activity during different stages of appetitive conditioning

To verify neuronal activity, we recorded calcium signals from NAc D1- and D2-neurons using the pyPhotometry recording setup (Akam and Walton, 2019) during appetitive Pavlovian learning.

We divided our analysis in early (Day1), intermediate (Day 6) and late (Day12) phases of learning. We also analyzed the differences in activity between trials in each of these days and no statistical differences were found for both D1- and D2-neuron activity (comparison between all trials, the first 5 trials, intermediate (middle) 5 trials and last 5 trials – Supplementary Figure 2A-R). This data led us to calculate the average of the activity of D1- or D2-neurons during the whole session.

Regarding day 1, D1-neurons showed a sustained increase in activity at CS onset (F_4,260_=1300, *p*<0.0001; [-2 to 0s] *vs* [0 to 2s], *p*<0.0001), which gradually decreased until US delivery (Figure 2A, B). Interestingly, D1-neuron net activity significantly decreases at US onset (F_6,260_=5087, *p*<0.0001; [-2 to 0s] *vs* [0 to 2s], *p*<0.0001; Figure 2A, B).

**Figure 2.**
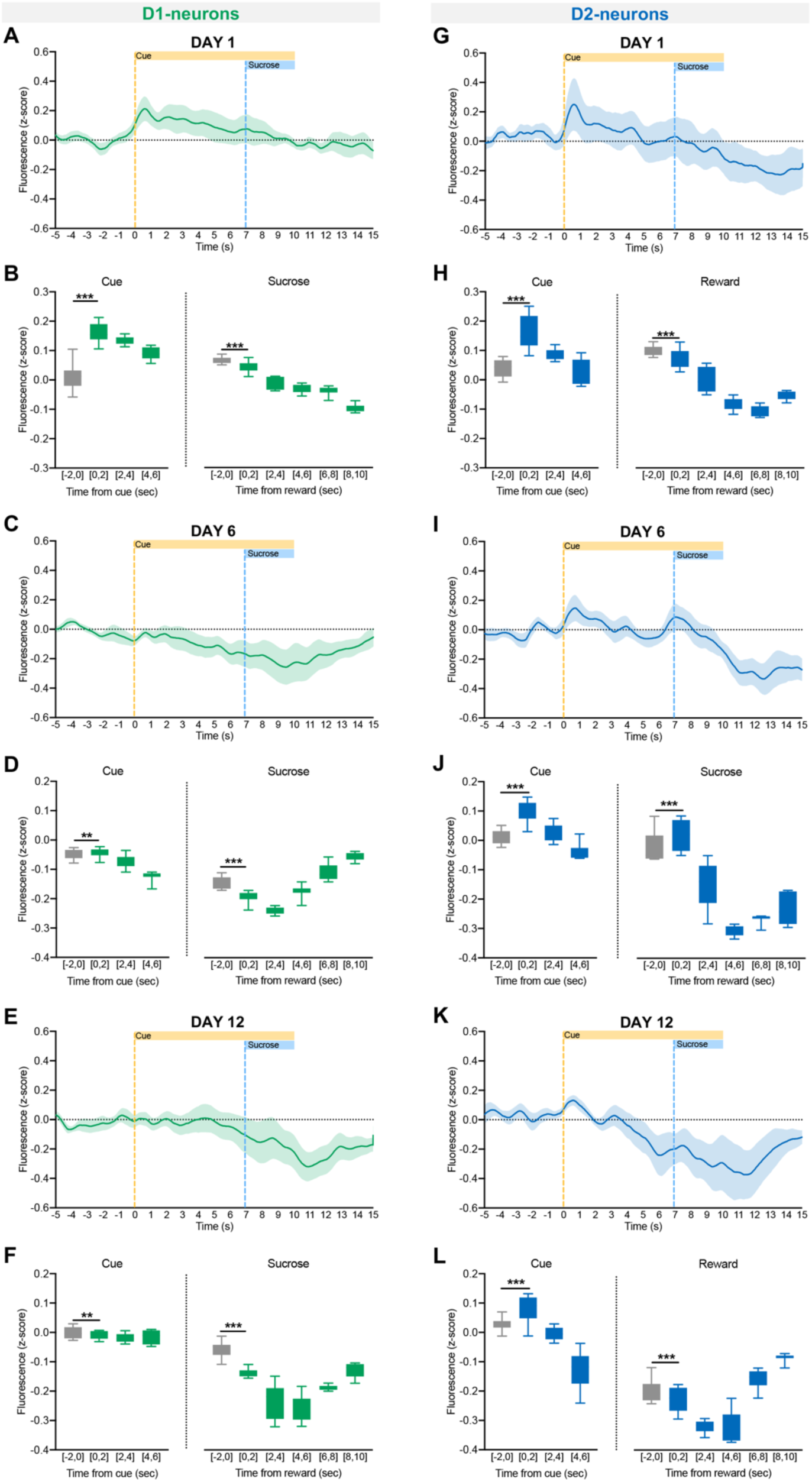
Dynamic activity of D1- and D2-neurons during different phases of appetitive Pavlovian conditioning. Z-score activity of D1-neurons in response to CS and US in the first (day 1 – **A**), intermediate (day 6 – **C**) and last session (day 12 – **E**) of appetitive learning. Box & Whiskers graphs showing mean activity of D1-neurons before and after CS or US exposure (day1 – **B**; day 6 – **D**; day 12 – **F**). Z-score activity of D2-neurons in response to CS and US in the **G** first, **I** intermediate and **K** last session of appetitive learning. Box & Whiskers graphs showing mean activity of D2-neurons before and after CS or US exposure (day1 – **H**; day 6 – **J**; day 12 – **L**). PSTH data are means ± SEM; Box & Whiskers data are min to max. ** p ≤ 0.01, *** p ≤ 0.001.

On day 6 of learning, we observe a slight transient increase in the activity of D1-neurons at CS onset (F_4,260_=4598, *p*<0.0001; [-2 to 0s] *vs* [0 to 2s], *p*=0.0017; Figure 2C, D). D1- neuron activity decreases after US onset in the PSTH (F_6,260_=2684, *p*<0.0001; [-2 to 0s] *vs* [0 to 2s], *p*<0.0001; Figure 2C, D).

On day 12, we observed a slight decrease in activity after CS onset in D1-neurons (F_4,260_=40.9, *p*<0.0001; [-2 to 0s] *vs* [0 to 2s], *p*=0.0035; Figure 2E, F). After US delivery, D1- neuronal activity robustly decreases (F_6,260_=1165, *p*<0.0001; [-2 to 0s] *vs* [0 to 2s], *p*<0.0001; Figure 2E, F).

Regarding D2-neurons, on day 1, we observe an increase in activity at CS onset (F_4,260_=1611, *p*<0.0001; [-2 to 0s] *vs* [0 to 2s], *p*<0.0001; Figure 2G, H), which gradually decreases and reaches baseline before US. After reward delivery, D2-neurons activity decreased for a prolonged period (F_6,260_=2980, *p*<0.0001; [-2 to 0s] *vs* all, *p*<0.0001; Figure 2G, H), up to 8 seconds after reward delivery (Figure 2G).

On day 6, we observed a transient increase in D2-neurons activity after CS onset (F_4,260_=1977, *p*<0.0001; [-2 to 0s] *vs* [0 to 2s], *p*<0.0001; Figure 2I, J). There was a transient increase in the activity during reward delivery (F_6,260_=1829, *p*<0.0001; [-2 to 0s] *vs* [0 to 2s], *p*<0.0001; Figure 2I, J), followed by a sharp decrease in activity that was sustained in time ([0 to 2s] *vs* [2 to 4s], *p*<0.0001; Figure 2I, J).

On day 12, when learning is fully established, D2-neuronal calcium activity increases at cue onset (F_4,260_=1384, *p*<0.0001; [-2 to 0s] *vs* [0 to 2s], *p*<0.0001; Figure 2K, L). There is a transient decrease in response to US ([-2 to 0s] *vs* [0 to 2s], *p*<0.0001; Figure 2K, L), and then D2-neurons maintain a decrease activity for 6 seconds, returning to baseline afterwards (Figure 2K, L).

In sum, these data suggest that the dynamic activity of D1- and D2-neurons is different throughout learning, particularly in the middle of the learning process. On day 1, there is an increase in activity in response to the CS in both populations, but this CS response at day 6 and 12 is only robustly present in D2-neurons. Both populations appear to decrease their activity in response to US as learning occurs, though the effect is more pronounced in D2-neurons in comparison to D1-neurons.

### D1- and D2-neuron responses to CS and US activity during appetitive Pavlovian conditioning

Considering the differences between D1- and D2-neurons as learning progresses, we evaluated the average calcium activity in response to CS and US onset over all days of the Pavlovian task. To do so, we calculated the average activity of each mouse to CS onset (from 0 to 2s; considering CS start at 0s) and to US onset (from 7 to 9s) across days of training.

Interestingly, the activity of D1-neurons decreases at CS period over days of training (Figure 3A), with a significant negative correlation between mean activity and days of training (r^2^=0.056, s=0.1790, *p*=0.0290; Figure 3A). The mean fluorescence is significantly lower at day 12 in comparison with day 1 (F_2,12_=6.9, *p*=0.035; day 1 *vs* day 12, *p*=0.0376; Figure 3B).

**Figure 3.**
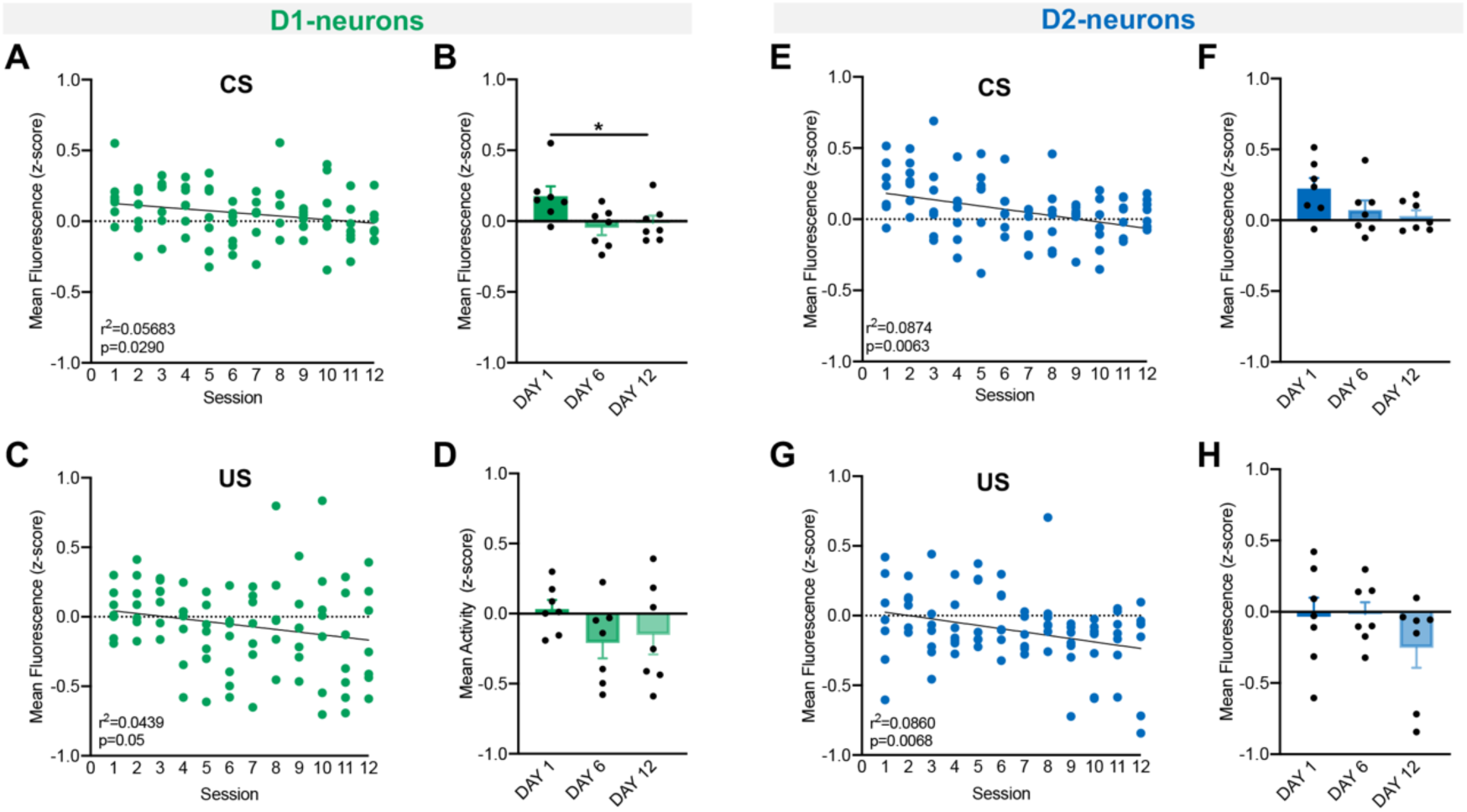
Temporal evolution of CS- and US-associated activity of D1- and D2-neurons during appetitive Pavlovian acquisition. Evolution of D1-neuronal activity during **A** CS and during **C** US across learning sessions. Quantification of mean fluorescence of D1-neurons in response to **B** CS and in response to **D** US across days of learning (comparison of day 1 vs day 6 vs day 12). Evolution of D2-neuronal activity during **E** CS and **G** US across learning sessions. Quantification of mean fluorescence of D2-neurons in response to **F** CS and **H** US across days of learning (comparison of day 1 vs day 6 vs day 12). Data are means ± SEM. * p≤0.05, ** p ≤ 0.01.

Regarding US period, we also observe a decrease in D1-neurons mean activity over days, observing a significant negative correlation (r^2^=0.043, s=0.312, *p*=0.05; Figure 3C). Despite this decrease, the mean fluorescence of D1-neurons was not statistically different on days 1, 6 and 12 (Figure 3D).

Regarding D2-neurons, we observed a significant negative correlation between mean activity at CS onset and days of training (r^2^=0.874, s=0.2533, *p*=0.0063; Figure 3E). Mean fluorescence over days was *quasi* significantly decreased (F_2,12_=4.2, *p*=0.0592; Figure 3F).

Regarding activity at US onset, a significant negative correlation between activity at US and days of training was present (r^2^=0.086, s=0.2697, *p*=0.0068; Figure 3G). However, no statistically significant differences were found in the analysis of the 3 days (Figure 3H).

In sum, these results show a gradual decrease in the activity of both D1- and D2- neurons in response to CS and to US throughout learning.

### Mice acquire proper responses to aversive Pavlovian conditioning

After the appetitive Pavlovian conditioning, the same mice (except for 1 D1-cre mouse that lost the implanted ferrule) performed an aversive Pavlovian conditioning paradigm (Figure 4A-I). Animals were subjected to a conditioning day, in which a CS (tone plus a cue light) was paired with a mild foot shock (US) of 0.5mA for 1 sec; 10 pairings were performed (Figure 4A). Following the conditioning session, mice were subjected to an omission session, in which the US (shock) was only present in 1/3 of the trials. On day 3, mice received a US extinction session, in which animals were subjected to 30 CS exposures with no US delivery.

**Figure 4.**
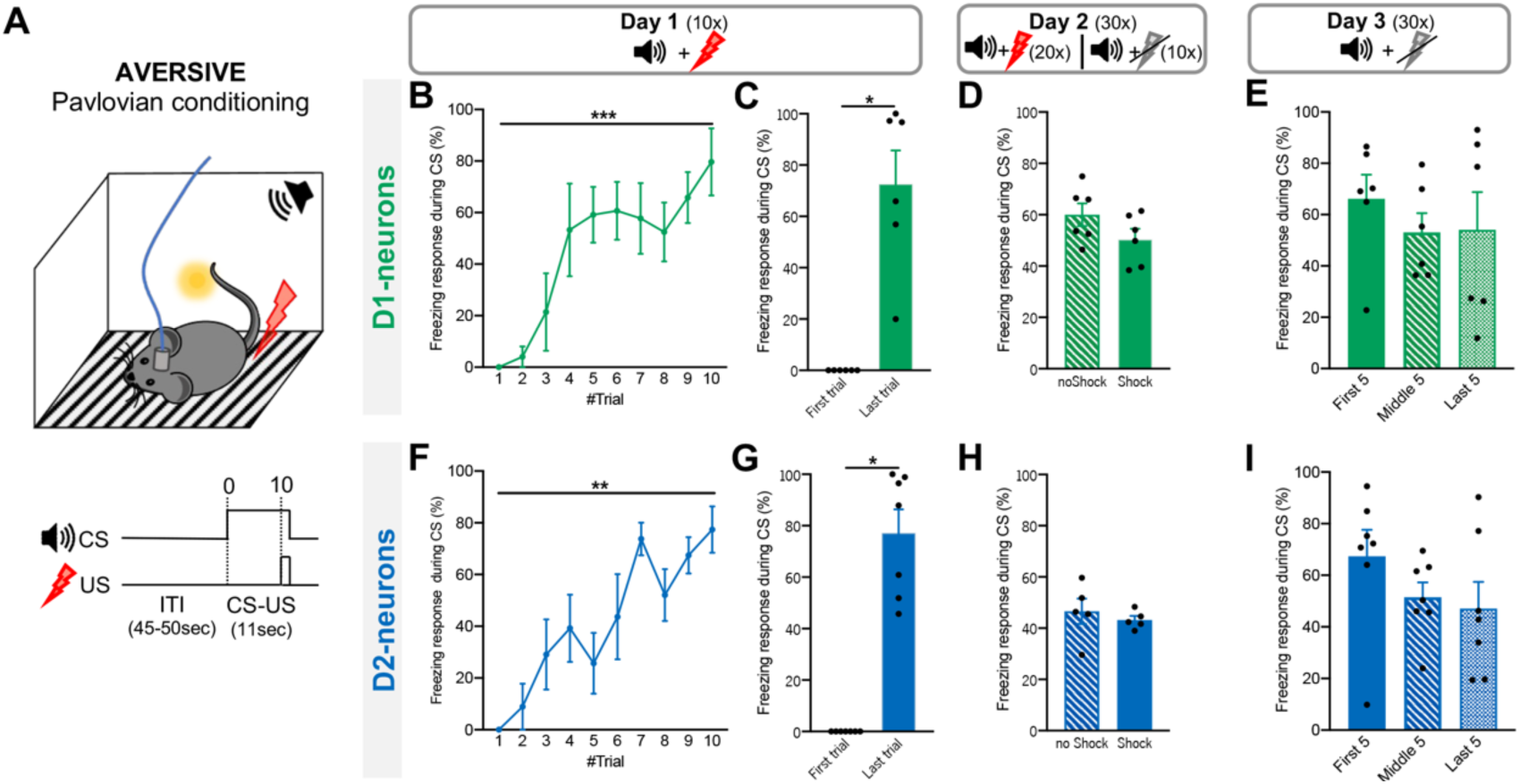
Acquisition of aversive Pavlovian conditioning by D1- and D2-cre mice during fiber photometry recordings of NAc neurons. **A** Aversive Pavlovian conditioning involved learning to associate a 3-kHz tone (CS) with a mild foot shock (US; 0.1mA, 1s). Freezing response in the aversive conditioning session of **B** D1-cre and **F** D2-cre mice. Freezing responses in the last trial were significantly higher than in the first trial in **C** D1-cre and **G** D2-cre mice. In the US omission session (1/3 of trials without US), freezing responses during CS were similar between Shock trials and noShock trials for **D** D1-cre and **H** D2-cre animals. In the US extinction trial, both **E** D1-cre and **I** D2-cre still present freezing responses to the CS. Data are means ± SEM. * p≤0.05, ** p ≤ 0.01, ***p ≤ 0.001.

On the conditioning session, we observe an increase in freezing over the trials for both D1-cre (F_10,6_=29.3, *p*=0.0006; Figure 4B) and D2-cre mice (F_10,7_=33.7, *p*=0.0001; Figure 4F). In addition, a significantly higher percentage of freezing was observed in the last trial when compared with the first trial (D1-cre: W_6_=21, *p*=0.0312, Figure 4C; D2-cre: W_7_=28, *p*=0.0156; Figure 4G). In the omission session, mice still presented a high percentage of freezing responses to the shock-predicting cue (D1-cre: Figure 4D; D2- cre: Figure 4H). All mice presented similar percentages of freezing in trials with shock delivery or shock omission, which is expected given the random occurrence of omission trials. In the US extinction session, both D1-cre (Figure 4E) and D2-cre (Figure 4I) mice presented freezing responses to the shock-predicting cue, despite absence of shock, even in the last trials (last 5 out of 30) of the session, indicating presence of CS-triggered fear memory.

### D1- and D2-neurons activity during aversive Pavlovian conditioning

During all sessions of the acquisition of the appetitive Pavlovian conditioning (Figure 4A) calcium transients were recorded and changes in activity were observed in both D1- and D2-neurons in response to the CS and US.

No significant differences in activity between trials were observed for both neuronal populations (comparisons in Supplementary Figure 5A-L). Thus, we averaged the activity of all trials.

On the conditioning day, D1-neurons responded to both CS and US. D1-neurons developed a transient increase in response to the CS (F_11,260_=1461, *p*<0.0001; [-2 to 0s] *vs* [0 to 2s], *p*<0.0001; Figure 5A, B). In response to US, D1-neurons developed a robust and sustained increase in activity ([8 to 10s] *vs* [10 to 12s], *p*<0.0001; Figure 5A, B), that only started to decrease three second after US onset ([12 to 14s] *vs* [14 to 16s], *p*<0.0001; Figure 5B).

**Figure 5.**
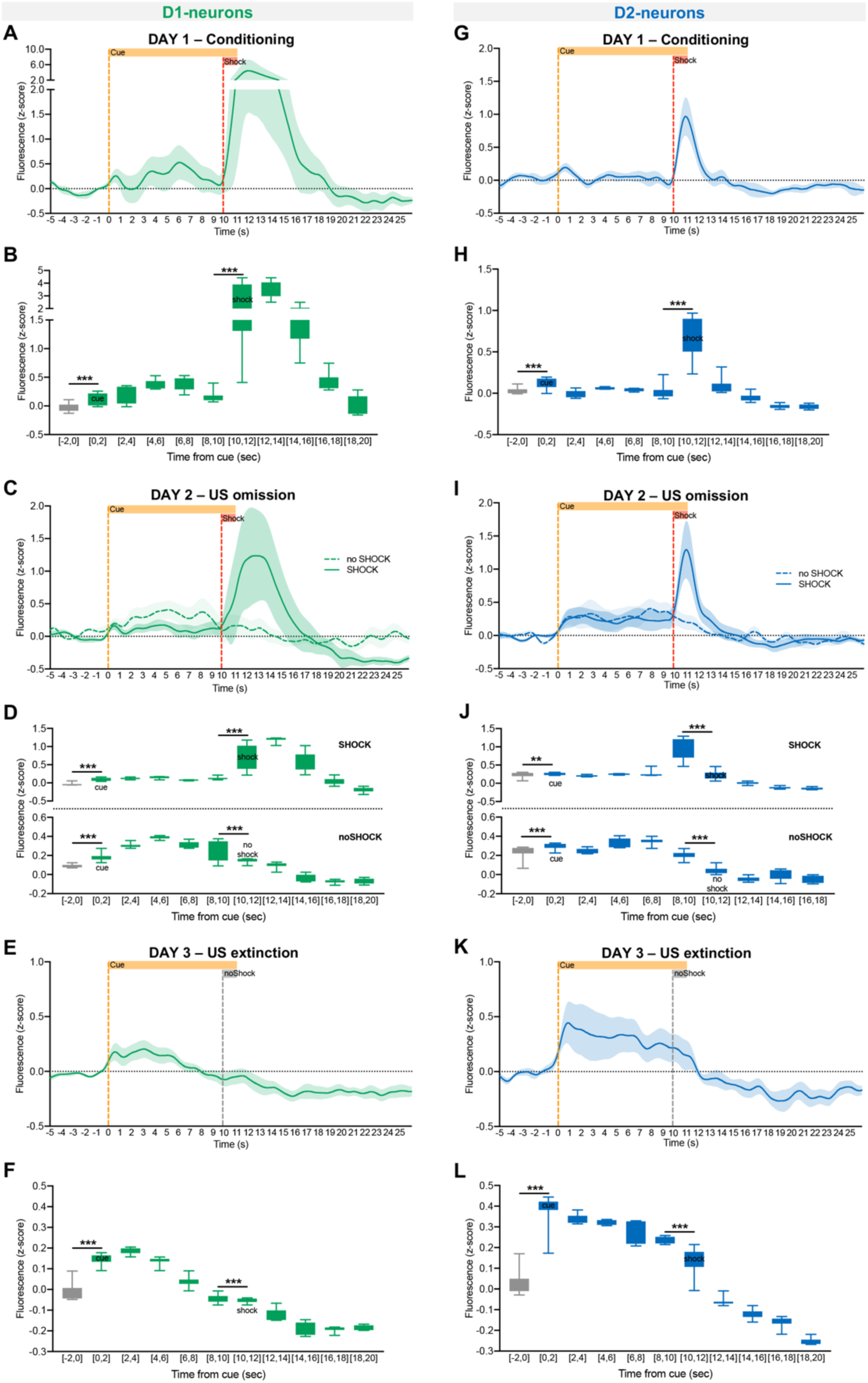
Dynamic activity of D1- and D2-neurons during aversive Pavlovian conditioning. **A** Z-score activity of D1-neurons in the acquisition session (cue-shock pairings), and **B** quantification of fluorescence before and after CS (cue) or US (Shock) exposure. **C** Activity of D1-neurons in the omission session (1/3 of cue-noShock pairings), and **D** quantification of fluorescence before and after cue or shock exposure, or cue exposure and noShock (omission). **E** Activity of D1-neurons in the extinction session (cue-noShock). **F** quantification of fluorescence before and after cue exposure and shock extinction. **G** Z-score activity of D2-neurons in the acquisition session, and **H** quantification of fluorescence before and after cue or shock exposure. **I** Activity of D2-neurons in the omission session, and **J** quantification of fluorescence before and after cue or shock exposure, or cue exposure and shock omission. **K** Activity of D2-neurons in the extinction session, and **L** quantification of fluorescence before and after cue exposure and shock extinction. PSTH data are means ± SEM; Box & Whiskers data are min to max. ** p ≤ 0.01, *** p ≤ 0.001.

D2-neurons showed a transient increase in activity in response to the cue (F_11,260_=2115, *p*<0.0001; [-2 to 0s] *vs* [0 to 2s], *p*<0.0001; Figure 5G, H), returning fast to baseline. In response to the US, D2-MSNs show a sharp significant increase in activity [8 to 10s] *vs* [10 to 12s], *p*<0.0001; Figure 5G, H), with the activity decreasing to baseline level 2 seconds after US exposure.

### D1- and D2-neurons respond to foot shock omission

On the day after conditioning, mice received 30 trials with 20 CS-US pairings and 10 trials of CS-only, randomly assigned (Figure 5J, M, Q).

On the CS-US trials, we observed similar activity to what was observed in the conditioning trials on day 1 (Figure 5C, D), with D1-neurons showing a subtle increase in activity at cue onset (F_11,260_=2387, *p*<0.0001; [-2 to 0s] *vs* [0 to 2s], *p*<0.0001; Figure 5C, D) that remained above baseline until US delivery. At US, D1-neuronal activity increased significantly in a sustained manner ([8 to 10s] *vs* [10 to 14s], *p*<0.0001; Figure 5C, D). On the CS only trials, D1-neurons increased activity in response to the CS (F_11,260_=4255, *p*<0.0001; [-2 to 0s] *vs* [0 to 2s], *p*<0.0001; Figure 5C, D) until the US, being then followed by a decrease to baseline levels after US omission ([8 to 10s] *vs* [10 to 12s], *p*<0.0001; Figure 5C, D).

Regarding D2-neurons, on day 2, a similar profile of activity was found as in day 1 for the CS-US trials, with a sustained increase in activity at CS onset (F_11,260_=2463, *p*<0.0001; [-2 to 0s] *vs* [0 to 12s], *p*<0.0001; Figure 5I, J). In response to US delivery, D2- neurons sharply increased activity ([8 to 10s] *vs* [10 to 12s], *p*<0.0001; Figure 5I, J). On the CS-only trials of day 2, D2-neurons presented a sustained increase in activity in response to CS (F_11,260_=4005, *p*<0.0001; [-2 to 0s] *vs* [0 to 10s], *p*<0.0001; Figure 5C), which only decreased after US omission ([8 to 10s] *vs* [10 to 12s], *p*<0.0001).

### D1- and D2-neuron activity in responses to foot shock extinction

On the 3^rd^ day, animals went through an extinction session, in which they were exposed to 30 trials of CS only. Calcium activity recorded during this session shows that both D1- and D2-neurons’ activity increases at CS onset (D1-neurons: F_11,260_=10975, *p*<0.0001; [-2 to 0s] *vs* [0 to 6s], *p*<0.0001, Figure 5E, F; D2-neurons: F_11,260_=9465, *p*<0.0001; [-2 to 0s] *vs* [0 to 10s], *p*<0.0001; Figure 5K, L). In response to shock omission, D1-neurons decrease activity ([8 to 10s] *vs* [10 to 20s], *p*<0.0001; Figure 5E, F), remaining low even after trial finish. Similarly, D2-neurons significantly decrease activity ([8 to 10s] *vs* [10 to 12s], *p*<0.0001), remaining below baseline for a long period of time.

In summary, these results show that D1-neurons present a more sustained response to the US period in comparison to D2-neurons that present a sharper CS response. On the other hand, in the US extinction day, D2-neurons show a prolonged increase in activity in response to CS.

## Discussion

Accumulating evidence has shown that the NAc plays an important role in mediating Pavlovian associations (Day et al., 2006; Day and Carelli, 2007; Roitman et al., 2005; Wan and Peoples, 2006), and that accumbal neurons respond to both CSs and USs (Roitman et al., 2005). However, there is still lack of information about the genetic nature of these neurons. Recent studies from our group have shown that optogenetic manipulation of both D1- and D2-MSNs can trigger positive and negative reinforcement (Soares-Cunha et al., 2020), depending on the pattern of MSN stimulation, suggesting that both populations can encode positive and negative associative learning. Here, we used fiber photometry to track bulk calcium transients in D1- and D2-neurons during appetitive and aversive Pavlovian conditioning.

Throughout appetitive Pavlovian training, D1- and D2-cre mice exhibit an increase in the number of nose pokes during CS presentation in relation to ITI, showing that animals learn the association between CS and US (sucrose). Importantly, this behavioral effect was accompanied by a gradual decrease in D1- and D2-neuron activity as animals learn the association between CS and US. This suggests that these two NAc subpopulations are dynamically changing activity as Pavlovian associations are acquired. Though there are studies showing that these subpopulations have an opposing function (Hikida et al., 2016, 2010; Kravitz et al., 2012; Volman et al., 2013), our data shows that both change activity during appetitive Pavlovian conditioning. In fact, these neurons appear to have a particularly relevant role in the early stages of cue-reward acquisition. There is an increase in the activity of D1-neurons during the CS period in the first day, but not on subsequent days of training. Conversely, D2-neurons present CS-triggered increase in activity over learning, suggesting different dynamics between the two neuronal populations during appetitive Pavlovian learning. Similarly, seminal electrophysiological studies have shown that NAc neurons develop responses to cues paired with sucrose and quinine (Roitman et al., 2005; Setlow et al., 2003).

Interestingly, both neuronal populations decrease activity during the US period across learning, in accordance with electrophysiological data showing that a large portion of NAc neurons decrease activity during reward delivery (Carelli, 2004; Janak et al., 2004; Roitman et al., 2005). In a recent study using calcium imaging recordings with miniaturized microscopes, it was found that reward consumption increased NAc lateral subregion activity but decreased medial subregion activity (Chen et al., 2023).

We also measured D1- and D2-neurons activity during an aversive Pavlovian conditioning in the same animals. As expected, animals exhibit robust freezing behavior in response to the cue in the last shock trial, in comparison to the first trial of conditioning. Moreover, in trials without shock (day 2 and day 3), animals still present freezing responses, showing that conditioning occurred.

Photometry data showed that both D1- and D2-neurons change activity during aversive Pavlovian conditioning. In response to US, D2-neuron response peaks were always sharper than D1-neurons. In agreement with our data, electrophysiological recordings in the NAc show that the delivery of quinine, an unpleasant taste stimulus, elicits mainly excitatory responses in NAc neurons (Roitman et al., 2005). In addition, increase in NAc firing rates was also observed in response to other aversive stimuli like air puff (Yanagimoto and Maeda, 2003).

Our data shows that D1- and D2-neurons also increase activity in response to CSs, in line with previous electrophysiological studies, though these studies did not identify the neuronal subtypes that were recording (Ray et al., 2022; Roitman et al., 2005; Setlow et al., 2003). Importantly, in US omission trials of day 2 and day 3, both populations decrease activity when the shock is omitted. This appears to signal US omission, which is quite an interesting finding that needs additional confirmation.

Our results corroborate a model of coexisting contribution of D1- and D2-MSNs for rewarding and aversive associative learning since both populations show significant activity changes. Yet, it is important to refer that fiber photometry calcium imaging reflects bulk population activity, lacking single-cell resolution. Considering recent single-cell RNA sequencing data, at least 18 MSN subtypes exist in the NAc (Chen et al., 2021), suggesting that even within D1- and D2-neurons, functional subpopulations can co-exist. Indeed, a study using miniaturized microscopes to monitor single cell activity in the NAc with calcium indicators in the medial and lateral shell shows distinct types of responses to unexpected rewards (water) in both D1- and D2-MSNs (Chen et al., 2023). This approach or electrophysiological recordings with optotagging of D1- or D2-neurons would be very interesting to perform in animals during appetitive and aversive Pavlovian conditioning to better understand the contribution of these neurons for these processes.

Overall, our results show a significant involvement of both NAc D1- and D2-neurons in reward and aversion processing, in line with previous optogenetic modulation data from our team (Soares-Cunha et al., 2020). Thus, this study reveals a more complex role played by accumbal D1- and D2-MSNs in encoding reward and aversion than the one proposed by the initial model of ventral basal ganglia function.

## Acknowledgements

CS-C and BC have Scientific Employment Stimulus contracts from the Portuguese Foundation for Science and Technology (FCT) (CEECIND/03887/2017; CEECIND/03898/2020). AVD and NG have FCT PhD grants (SFRH/BD/147066/2019; 2022.12973.BD).

This work received funding from the Bial Foundation grant (175/2020). In addition, part of the experiments was funded by the European Research Council (ERC) under the European Union’s Horizon 2020 research and innovation programme (grant agreement No 101003187) and by the “la Caixa” Foundation (ID 100010434), under the agreement LCF/PR/HR20/52400020. Part of the work received funding from Portuguese Foundation for Science and Technology (FCT) under the scope of the projects PTDC/MED-NEU/4804/2020, PTDC/SAU-TOX/6802/2020, 2022.02201.PTDC and 2022.01467.PTDC.

This work was also supported by a FEBS (Federation of European Biochemical Societies) Excellence Award (2022) and IBRO Early Career Award (2022), attributed to CS-C.

Part of this work also received funding from the ICVS Scientific Microscopy Platform, member of the national infrastructure PPBI - Portuguese Platform of Bioimaging (PPBI-POCI-01-0145-FEDER-022122); and by National funds, through the FCT - UIDB/50026/2020 and UIDP/50026/2020.

## Contributions

Conceptualization, CD, AVD, CSC, and AJR; investigation, CD, AVD, TC, BC, LP, NS, NVG; data analyses, CD, AVD, TC and RFO; statistical analyses, CD, TC and CS-C; writing - original draft preparation, CD and AVD; writing, CSC and AJR; review and final proofing of the manuscript – all authors.

## Corresponding authors

Correspondence to Carina Soares-Cunha or Ana João Rodrigues

---

**Supplementary Figure 1.**
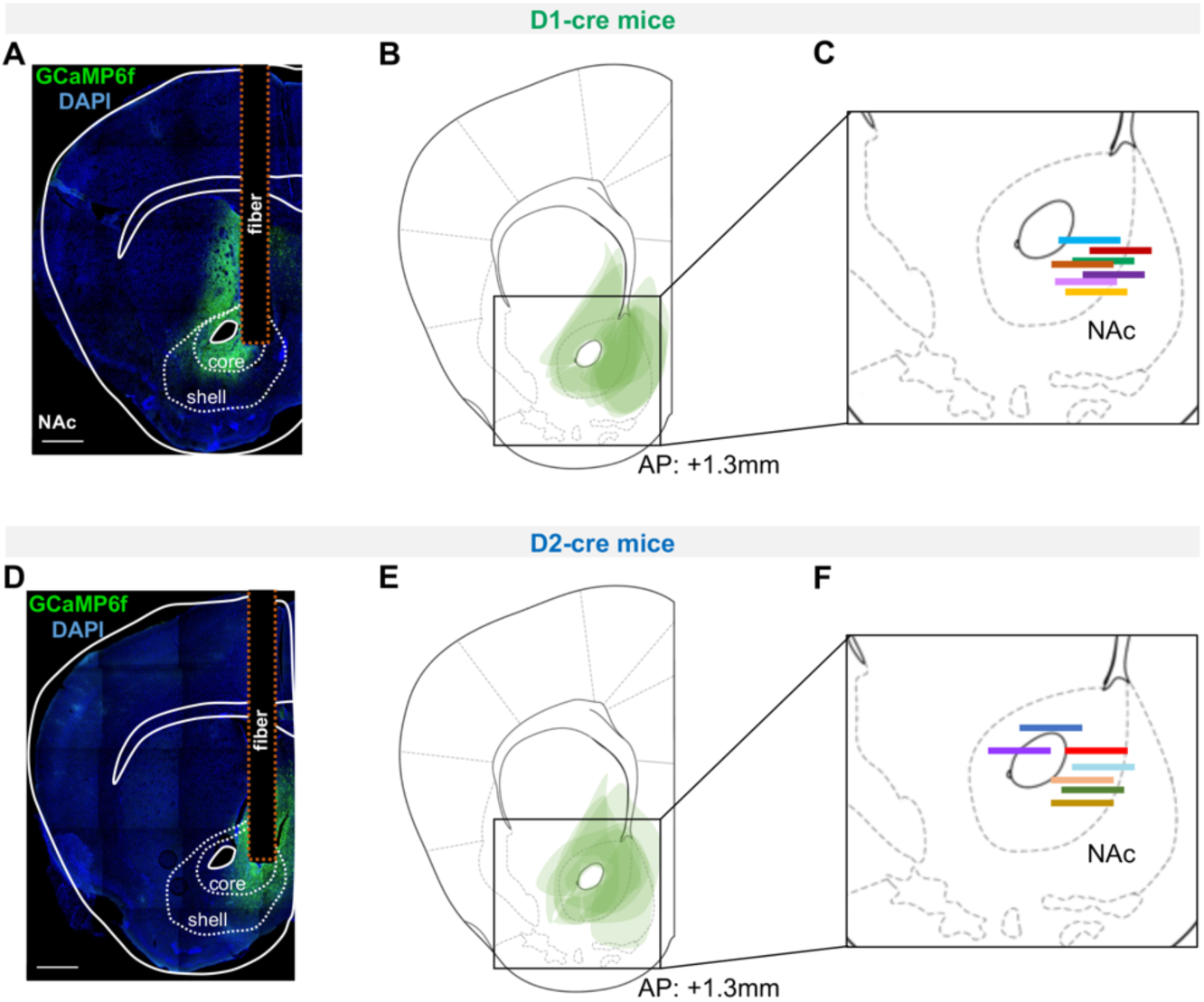
Viral expression and optic fiber placement in the NAc of mice used for fiber photometry recordings during appetitive and aversive Pavlovian conditioning. **A, D** Coronal slice showing expression of GCaMP6f in the NAc of a D1-cre and a D2-cre mouse, respectively, using immunostaining for YFP; scale bar: 500 μm. **B, E** Representation of viral expression spreading in the NAc of D1-cre and D2-cre mice, respectively. **C, F** Location of the tip of the implanted optic fiber in D1-cre and D2-cre mice, respectively. n_D1-cre_=7, n_D2-cre_=7.

**Supplementary Figure 2.**
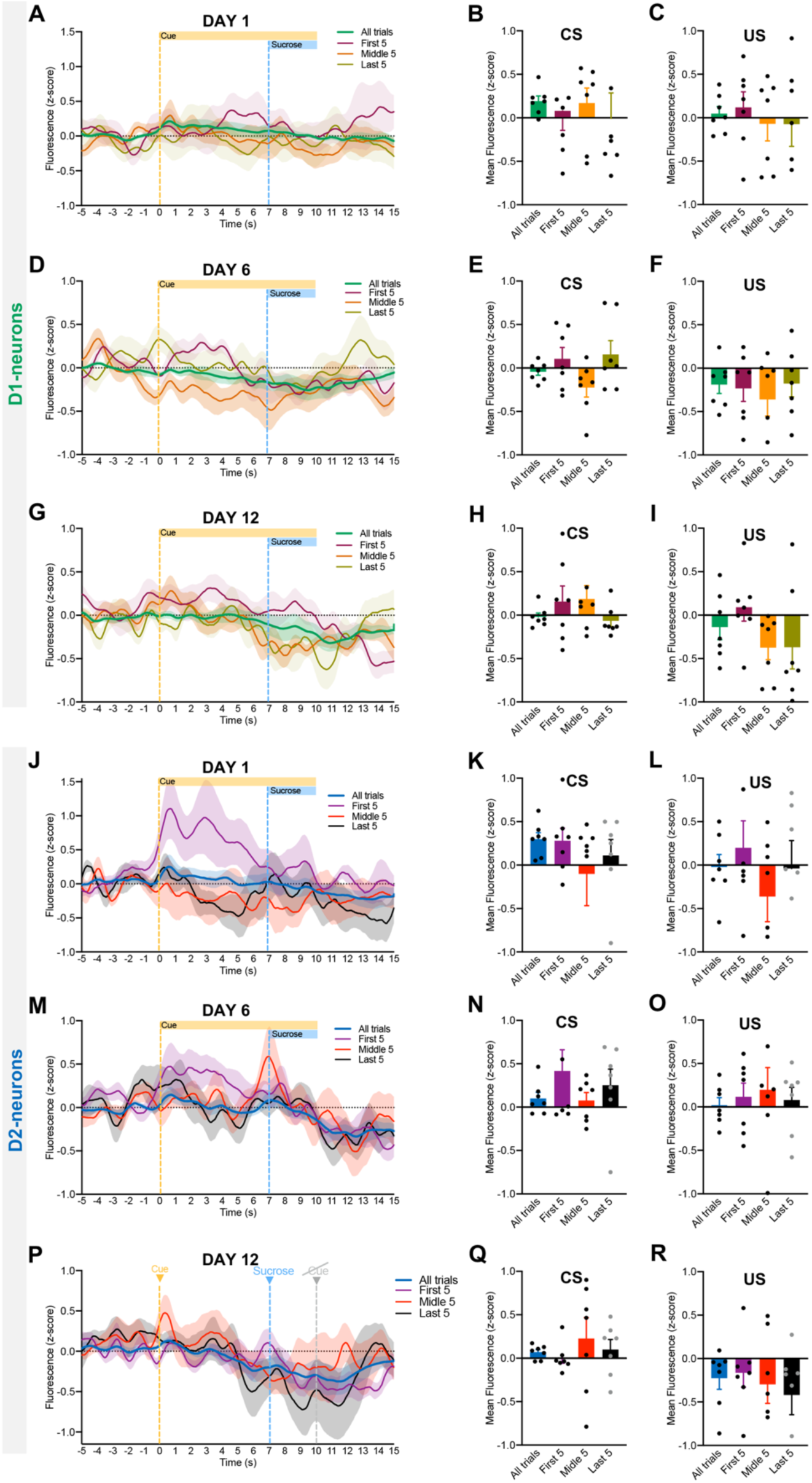
Dynamic activity of D1- and D2-neurons during different phases of appetitive Pavlovian learning. Z-score activity of D1-neurons in response to CS and US in the first (day 1 – **A**), intermediate (day 6 – **D**) and last session (day 12 – **G**) of appetitive learning, separated by the first 5, intermediate 5 and last 5 trials. Bar graphs showing mean activity of D1-neurons before and after CS or US exposure in the first 5, intermediate 5 and last 5 trials of each session (day1 CS and US – **B, C,** respectively; day 6 CS and US – **E, F,** respectively; day 12 CS and US – **H, I,** respectively). Z-score activity of D2-neurons in response to CS and US in the first (day 1 – **J**), intermediate (day 6 – **M**) and last session (day 12 – **P**) of appetitive learning, separated by the first 5, intermediate 5 and last 5 trials. Bar graphs showing mean activity of D2-neurons before and after CS or US exposure in the first 5, intermediate 5 and last 5 trials of each session (day 1 CS and US – **K, L,** respectively; day 6 CS and US – **N, O,** respectively; day 12 CS and US – **Q, R,** respectively). Data are means ± SEM.

**Supplementary Figure 3.**
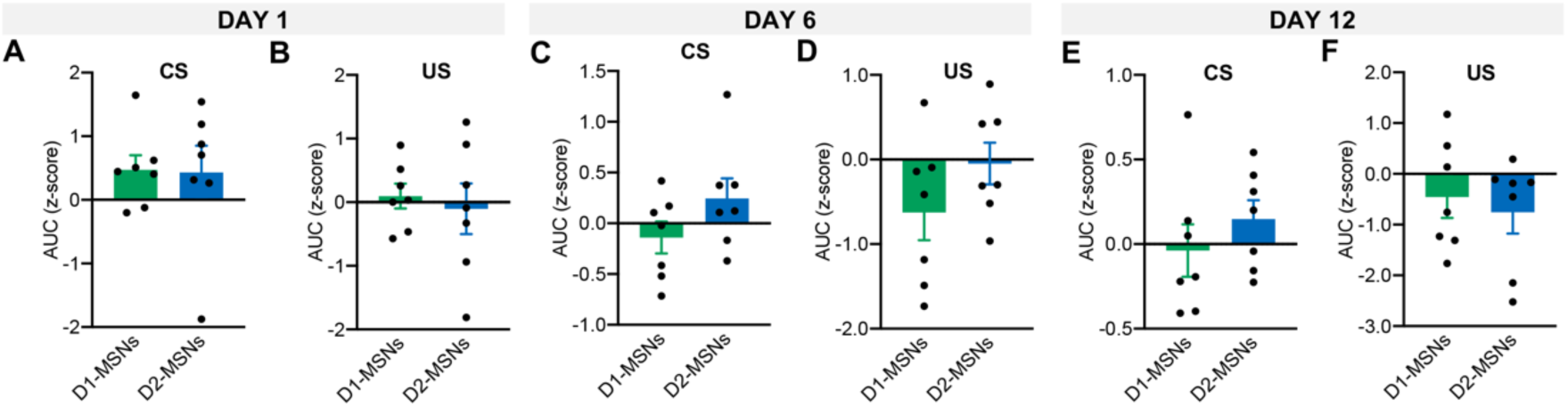
Comparison of D1- and D2-neurons activity during different phases of appetitive Pavlovian learning. Bar graphs showing area under the curve (AUC) of D1- and D2-neurons during **A** CS or **B** US exposure on day 1 of learning. Bar graphs showing AUC of D1- and D2-neurons during **C** CS (U_34,71_=6, *p*=0.0175) or **D** US exposure on day 6 of learning. Bar graphs showing AUC of D1- and D2-neurons during **E** CS or **F** US exposure on day 12 of learning. Data are means ± SEM.

**Supplementary Figure 4.**
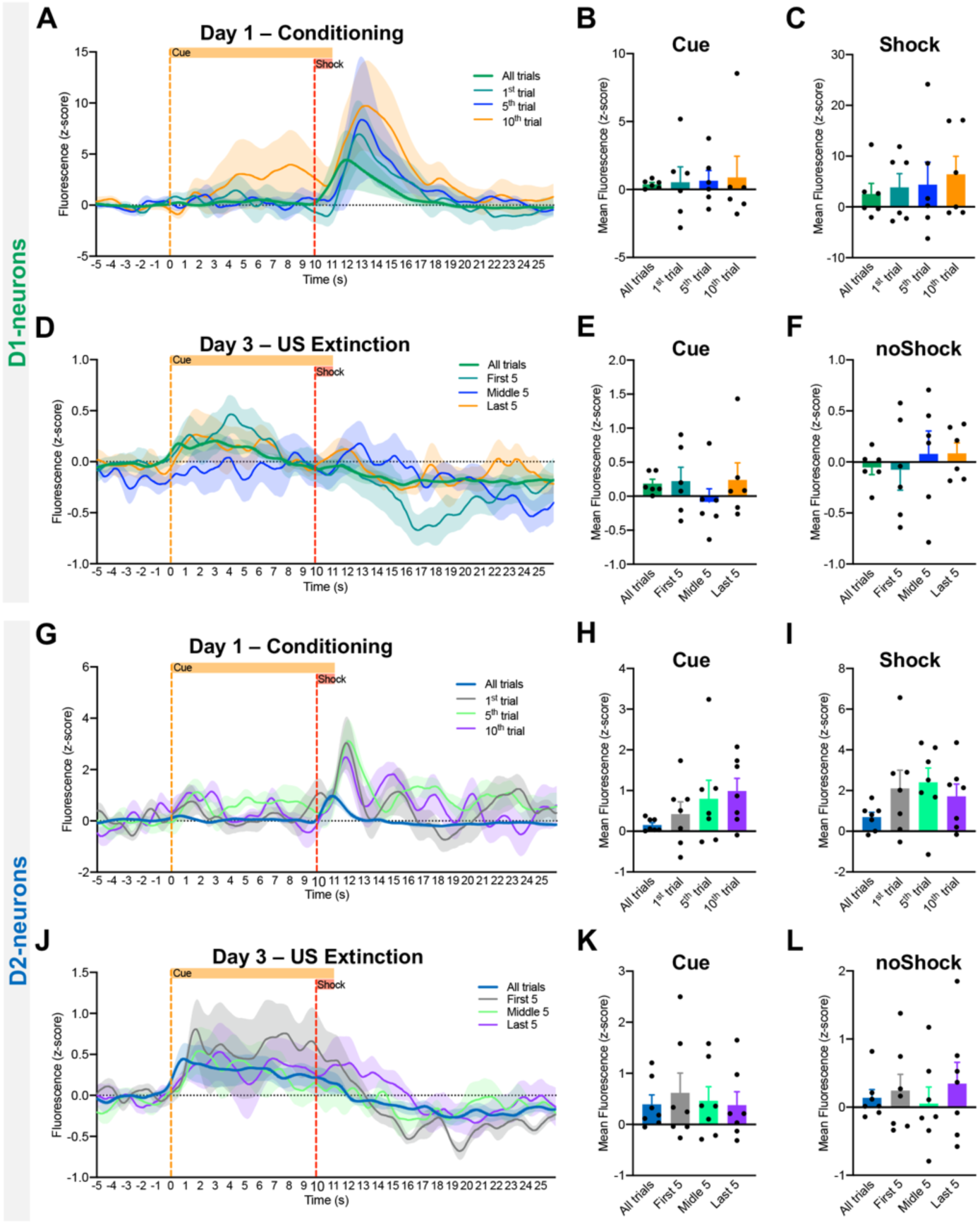
Dynamic activity of D1- and D2-neurons during different phases of aversive Pavlovian learning. **A** Z-score activity of D1-neurons in response to cue and shock in the conditioning session of aversive learning, separated by the first 5, intermediate 5 and last 5 trials. Bar graphs showing mean activity of D1-neurons before and after cue or shock exposure in the first 5, intermediate 5 and last 5 trials of each session (cue and shock – **B, C,** respectively). **D** Z-score activity of D1-neurons in response to cue exposure and shock extinction in the extinction session of aversive learning, separated by the first, fifth and tenth trial. Bar graphs showing mean activity of D1-neurons before and after cue exposure or shock extinction exposure in the first, fifth and tenth trial (cue and noShock – **E, F,** respectively). **G** Z-score activity of D2-neurons in response to cue and shock in the conditioning session of aversive learning, separated by the first 5, intermediate 5 and last 5 trials. Bar graphs showing mean activity of D2-neurons before and after cue or shock exposure in the first 5, intermediate 5 and last 5 trials of each session (cue and shock – **H, I,** respectively). **J** Dynamic activity (z-score) of D2-neurons in response to cue exposure and shock extinction in the extinction session of aversive learning, separated by the first, fifth and tenth trial. Bar graphs showing mean activity of D2-neurons before and after cue exposure or shock extinction exposure in the first, fifth and tenth trial (cue and noShock – **K, L,** respectively). Data are means ± SEM.

**Supplementary Figure 5.**
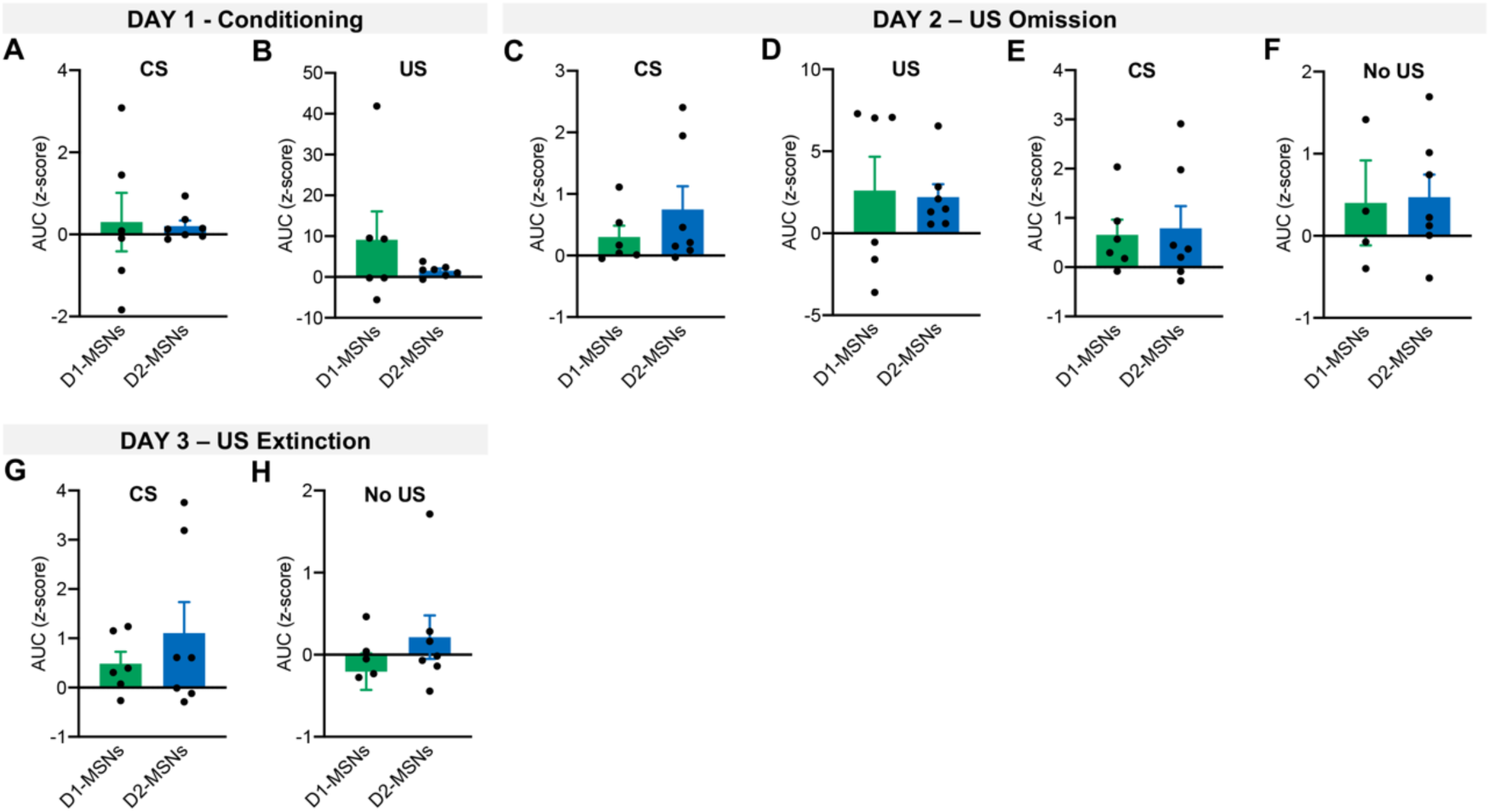
Comparison of D1- and D2-neurons activity during different phases of aversive Pavlovian learning. Bar graphs showing AUC of D1- and D2-neurons during **A** CS or **B** US exposure on the conditioning session. Bar graphs showing AUC of D1- and D2-neurons during **C** CS or **D** US exposure on the CS-US trials of US omission session. Bar graphs showing AUC of D1- and D2-neurons during **E** CS or **F** no-US exposure on the CS only trials of US omission session. Bar graphs showing AUC of D1- and D2-neurons during **G** CS or **H** no-US exposure on the US extinction session. Data are means ± SEM.

